# ZFP36 is a post-transcriptional regulator of the P2X4 receptor in myeloid cells

**DOI:** 10.64898/2026.10.06.757012

**Authors:** Damien Gilabert, Negar Ahmadi, Lucie Gentil, Valentine Charmes, Nathalie Linck, François Rassendren, Lauriane Ulmann

## Abstract

P2X4 is an ATP-gated purinergic receptor upregulated in spinal microglia after peripheral nerve injury, where it plays a key role in neuropathic pain. Yet, the molecular mechanisms controlling P2X4 expression remain incompletely understood.

To address this question, we examined P2X4 mRNA expression and translation in spinal microglia following nerve injury. While total P2X4 mRNA levels remained unchanged, ribosome-associated P2X4 mRNA increased, revealing a previously unrecognized level of translational regulation. We therefore investigated whether the RNA-binding protein ZFP36 (tristetraprolin), a well-established regulator of inflammatory mRNAs, controls P2X4 expression. Our results demonstrate that the P2X4 3′UTR strongly repressed reporter and protein expression and that ZFP36 emerged as a negative regulator of P2X4 mRNA. This regulation involved phosphorylation-dependent signaling through p38 MAPK and 14-3-3. Moreover, ZFP36 knockdown increased P2X4 protein expression, while manipulation of ZFP36 phosphorylation altered P2X4-dependent ATP-evoked calcium responses.

Together, these findings identify ZFP36 as a previously unrecognized post-transcriptional regulator of P2X4, extending its role beyond inflammatory mediators to a functional purinergic receptor. They reveal post-transcriptional control of receptor availability as a novel regulatory layer linking myeloid cell state to purinergic responsiveness and provide a new framework for understanding microglial P2X4 regulation in neuropathic pain.

## Introduction

Extracellular adenosine triphosphate (ATP) is a crucial signaling molecule that modulates neuro-immune communication via its action on purinergic receptors, encompassing the ligand-gated P2X receptor family [1][2]. Among them, P2X4 is expressed in microglia and immune cells. When activated by ATP, calcium influx guides signaling pathways involved in inflammatory responses, the release of cytokines and the control of neuronal activity [3][4]. Growing evidence indicates that P2X4 plays a particularly important role in pathological conditions linked to neuroinflammation. After peripheral nerve injury, P2X4 expression is markedly increased in the dorsal horn of the spinal cord microglia, leading to the release of brain-derived neurotrophic factor (BDNF) and subsequent modifications in neuronal chloride homeostasis, dorsal horn neurons hyperexcitability, thereby contributing to neuropathic pain hypersensitivity. In addition to the nervous system, P2X4 receptors regulate macrophage inflammatory signaling pathways, indicating that precise control of P2X4 expression is crucial for maintaining immune homeostasis [5]. Although the physiological and pathological relevance of P2X4 is increasingly acknowledged, the molecular mechanisms governing its expression and regulation remain incompletely understood.

Post-transcriptional regulation of mRNA stability represents a major mechanism controlling gene expression. RNA-binding proteins (RBPs) that recognize AU-rich elements (AREs) in the 3′ untranslated regions (3′UTRs) of transcripts rapidly modulate mRNA turnover and translation in response to cellular stimuli. ZFP36, also known as tristetraprolin (TTP), promotes the degradation of ARE-containing transcripts [6]. ZFP36 plays a central role in limiting inflammatory responses by destabilizing mRNAs encoding cytokines and other immune mediators, including tumor necrosis factor-α (TNF-α) [7]. Consistent with this function, genetic deletion of *Zfp36* in mice results in a severe systemic inflammatory syndrome driven by uncontrolled cytokine production [8]. Further studies have highlighted the importance of ZFP36 in immune cells, showing that its disruption in T cells accelerates their response and enhances the production of key pro-inflammatory cytokines such as IFN-γ and TNF-α, which can improve antiviral immunity [9].

Although both purinergic signaling and ARE-mediated mRNA decay are key regulatory layers in inflammatory pathways, potential connections between these mechanisms remain poorly characterized. Increasing evidence indicates that RNA-binding proteins can directly regulate the expression of receptors and signaling molecules involved in immune responses, thereby shaping cellular sensitivity to extracellular cues [10]. The roles of ZFP36 family proteins in inflammatory diseases, with genetic variants being associated with autoimmune diseases such as rheumatoid arthritis, multiple sclerosis, and lupus, suggest that dysregulation of these RBPs contributes to chronic inflammation in humans [11][12]. This raises the possibility that post-transcriptional regulators such as ZFP36 may influence purinergic signaling by controlling the expression or function of receptors such as P2X4.

Here, we provide evidence that ZFP36 regulates P2X4 expression through AU-rich regulatory elements within the P2X4 3′UTR, thereby limiting receptor expression and function. Under neuropathic conditions, increased ribosome-associated P2X4 mRNA is consistent with enhanced translation in spinal microglia. Collectively, our findings reveal a post-transcriptional mechanism that shapes purinergic responses and identify ZFP36 as a candidate regulator of P2X4-driven neuroinflammatory processes.

## Methods

### Mice

C57BL/6J and *Cx3Cr1*^CreERT2/+^;*Rpl22*^HA(RiboTag)/+^ female and male mice from 6 to 10 weeks old were housed under a standard 12 h light/dark cycle with food and water available ad libitum. Mice were bred in the SPF animal facility of the Institute for Functional Genomic (IGF, Montpellier, France; Agreement from the Ministry of Agriculture N° D34-172-13). All experimental procedures complied with official European guidelines for the care and use of laboratory animals (Directive 2010/63/UE) and were approved by the ethical committee of Montpellier (CEEA36) and the French Ministry of Agriculture APAFIS#22741– 2019102914341098 v4).

### Spared nerve injury (SNI)

Mice were randomly assigned to SNI or sham group. Animals were anesthetized under 3% isoflurane/oxygen, the left sciatic nerve was exposed and the peroneal and tibial branches were ligated and transected. Skin was closed with silk sutures. An identical procedure was performed in sham-operated mice except the nerve ligation and transection. The mice were isolated until they had fully recovered from the surgery before being reintroduced with their littermates.

### Tamoxifen

100 µl of Tamoxifen (10mg/mL, T5648, Sigma) was injected intraperitoneally for 3 consecutive days. Mice were used 21 days post injection, allowing for peripheral CX3CR1 +cells replenishment while microglia remained depleted.

### Tissue preparation

Mice were euthanized with Euthasol Vet® (300 mg/kg) and perfused with PBS. Spinal cords were harvested and placed overnight in 4% paraformaldehyde. The contralateral side of the lesion was recognized by a scalpel’s notch. The L1 to S1 segments were cut with a vibratome (VT1000S, Leica) into 40 μm sections and stored in PBS +0.1% sodium azide at 4°C. When indicated, lumbar segments were cryoprotected in a solution of 20% sucrose within 0.1M PBS overnight and stored at −80°C. Using cryostat (Leica), series of 50 μm-thin coronal slices of the spinal cord were obtained.

### Immunohistochemistry

The sections were washed with PBS. Non-specific sites were saturated with 10% donkey serum (S30, Millipore) diluted in PBS +0.3% Triton X-100 (Sigma, X100) for 1h at room temperature. Primary antibody (rabbit anti-Iba1, 1:2000, AB178846, Abcam), was incubated overnight at 4°C in DPBS +0.3% Triton X-100. After three washes in PBS, the sections were incubated for 2h at room temperature with the corresponding secondary antibody (donkey anti-rabbit CF488A, 1:1000, SAB4600036, Sigma) diluted in PBS +0.3% Triton X-100. The sections were rinsed with PBS 3 × 10 min and then mounted between slide and coverslip with mounting medium (Dako, S3023). L4 to L6 sections were observed using a fluorescence microscope (Axioimager 2, Zeiss) with an apotome grid module (Apotome 3, Zeiss) equipped with a digital camera (cool-snap HQ, PhotoMetrics) or a fluorescence confocal microscope (SP8-UV, Leica).

### Microglial RiboTag immunoprecipitation and RNA extraction

Tamoxifen-treated *Cx3Cr1*^CreERT2/+^;*Rpl22*^HA(RiboTag)/+^ mice were used 15 days after surgery. Animals were euthanised with Euthasol Vet® (300 mg/kg, i.p.) and perfused intracardially with DPBS. The lumbar spinal cord was cut lengthwise into ipsilateral/contralateral and dorsal/ventral quarters and snap-frozen at −80 °C. A thoracic segment was immersion-fixed in 4 % PFA as a recombination control. Quarters were pooled in sets of four and homogenized (Potter-Elvehjem) in 1 ml lysis buffer (1 % NP-40, 100 mM KCl, 50 mM Tris pH 7.4, 12 mM MgCl₂, 1 mg/ml cycloheximide, 10 mg/ml heparin, 10 mM RNaseOUT (Invitrogen, Thermo Fisher Scientific), 50 mM DTT, protease inhibitors). A 100-µl aliquot was stored at −80 °C as total-RNA input. The remaining lysate was incubated overnight at 4 °C with an anti-HA antibody (16B12, BioLegend), then overnight at 4 °C with protein G–coupled magnetic beads (Dynabeads™, 10004D, Thermo Fisher Scientific) pre-blocked twice in lysis buffer and recovered on a magnetic rack (Invitrogen™ DynaMag™, 12321D, Thermo Fisher Scientific). Beads were washed twice in high-salt buffer (300 mM KCl, 1 % NP-40, 50 mM Tris pH 7.4, 12 mM MgCl₂, 10 µg/ml cycloheximide, 1 mM DTT) and eluted in RLT buffer (Qiagen RNeasy, 79216) containing 1 % β-mercaptoethanol.

RNA was purified with the RNeasy Plus Mini kit (Qiagen, 74136) for total RNA or the RNeasy Micro kit (Qiagen, 74004) for microglial RNA, including on-column DNase I digestion (Qiagen, 79254; 15 min, room temperature), and eluted twice in RNase-free water. Total RNA was quantified on a NanoDrop 2000 (Thermo Fisher Scientific); immunoprecipitated RNA was quantified and quality-checked on a 2100 Bioanalyzer (Agilent), all samples showing RIN > 7. cDNA was synthesized with the iScript reverse transcription kit (Bio-Rad, 1708891BUN) and diluted five-fold.

### Droplet digital PCR

ddPCR was performed on the QX200 Droplet Digital PCR system (Bio-Rad) with ddPCR Supermix for Probes (no dUTP; Bio-Rad, 1863025) according to the manufacturer’s instructions. Reactions (22 µl) contained supermix, cDNA, and primers and probes at 600 nM and 250 nM, respectively (*Tmed10*: forward CAGGTGGATGGGAAAGATGTA, reverse GCCATGCAAGAGGAATTTATG, probe AGGCTGTGCTGTTAAGGTTCCAGT; *P2rx4*: forward CCAACACTTCTCAGCTTGGAT, reverse TGGTCATGATGAAGAGGGAGT, probe TATGTGGTCCCAGCTCAGGAGGA). Twenty microliters were partitioned into ∼20 000 droplets on a QX200 droplet generator (Bio-Rad, 1814000), giving a Poisson distribution of cDNA molecules, and amplified in a sealed 96-well plate on a Mastercycler ep gradient thermal cycler (Eppendorf). The lid was heated at 105 °C and the PCR program was: 95 °C for 10 min, 40 cycles of 95 °C for 30 s and 55 °C for 1 min (ramp 2.5 °C/s), then 98 °C for 10 min. Droplets were read on a QX200 reader (Bio-Rad; FAM and HEX channels) and analyzed in QuantaSoft (Bio-Rad).

### Western blot

Cells were homogenized in lysis buffer (20 mM Hepes, pH 7.4, 100 mM NaCl, 5 mM EDTA, 1% Triton X-100 and Complete Protease Inhibitor cocktail (Roche)). Lysates were clarified by centrifugation and protein concentration was determined using a protein assay kit (Bio-Rad). Proteins were separated on 4-12% NuPage precast gel (Life technologies), and transferred onto nitrocellulose membranes. The membrane was blocked with 5% nonfat dry milk/0.5% Tween20 in phosphate buffer saline (PBST) for 3 h at room temperature. The membrane was then incubated overnight at 4°C with the appropriate antibody diluted in PBST: rabbit anti-P2X4 (1/1000, Sigma) rabbit anti-GFP (1/1000, Torrey Pines), rabbit anti-ZFP36 (1/500, Cell Signaling) and mouse anti-tubulin (1/1000, Sigma). After three washes in PBST, the membrane was incubated with specific HRP-conjugated secondary antibody for 1 h at room temperature and revealed with ECL1 detection kit (GE Healthcare). Images were acquired with a Chemidoc Touch (Biorad) and analyzed with ImageLab software.

### RBPmap analysis of the P2X4 3’ UTR

To identify potential RNA binding site in mouse P2X4 3’ UTR sequence, we us the web-based tool RBPmap (http://rbpmap.technion.ac.il/index.html). RBPmap is a computational tool that enables accurate prediction and mapping of RNA binding proteins (RBPs) binding sites on any RNA sequence [13]. The 1424 bp 3’UTR sequence of mouse P2X4 was used as the query sequence, using high stringency and conservation-based filters. Among all RNA binding proteins identified as potentially binding to the UTR, ZFP36 was selected based on its strong expression in myeloid cells and its known role in the regulation of cytokine expression.

### Cell culture

Bone marrow cells were obtained by flushing femurs and tibias with sterile DMEM using a 25G needle. The suspension was centrifuged at 300 × g for 5 minutes and the pellet was directly resuspended in DMEM supplemented with 10% FBS and 30% L929 cell-conditioned medium. Cells were seeded in non-treated 10 cm Petri dishes and incubated at 37 °C with 5% CO₂. On day 3, half of the medium was replaced with fresh supplemented medium. Cells were considered fully differentiated into macrophages on day 7 and used for downstream experiments. Primary microglial cells were cultured as previously described [4].

HEK cells were grown in DMEM supplemented with 10% fetal heat-inactivated bovine serum (FBS) and 1% penicillin/streptomycin. Plasmids were transfected using lipofectAMINE 2000 (Invitrogen) according to the manufacturer’s instructions. Cells were incubated overnight and were passaged the following day and lysates were used on day 2 post-transfection.

### Calcium imaging

HEK cells were transfected with pTRIPZ Myc2-mouse TTP WT and pTRIPZ Myc2-mouse TTP aa S52 S178 MUT plasmids and plated in 96-well plates and loaded with 1.25 µM Cal-520®, AM (21130, AAT Bioquest) in calcium buffer (135 mM NaCl, 5 mM KCL, 2 mM CaCl2, 2 mM MgCl2, 10 mM glucose, 10 mM Hepes, pH 7.4) for 1h at 37°C. After washing, cells were stimulated with ATP (20 µM) and signals were detected at 480/540 nm with the FDSS®/μCell (Functional Drug Screening System, Hamamatsu).

Primary macrophage transduction One week after plating, BMDMs were rinsed with phosphate-buffered saline (PBS) and treated with Versene to promote cell detachment. Cells were subsequently replated in 6-well plates. Two days after replating, cells were transduced with 5 µL of pWPT-EF1α-GCaMP6s-P2A-Scarlet lentiviral vector together with either 10 µL of the pTRIPZ-Myc-ZFP36 or 10 µL of the pTRIPZ-Myc-ZFP36-mut lentiviral vector. To enhance transduction efficiency, 5 µL of LentiBlast (LBPX1500, Oz Biosciences) was added per well. Transduced cells were selected with puromycin (3 µg/mL; HB4041, HelloBio) for two days at DIV11 and DIV14. Following selection, cells were transferred to imaging dishes (81156, ibidi). Four hours after replating, doxycycline (100 ng/mL; D9891, Sigma-Aldrich) was added to induce transgene expression for 24 h prior to imaging experiments

### Live-cell fluorescence imaging

Live-cell fluorescence imaging experiments were performed using an Olympus IX70 epifluorescence microscope equipped with a ×20/1.35 objective, a pE-800 LED illumination system (CoolLED), and an Evolve 512 EMCCD camera (Photometrics) controlled by MetaFluor software (Molecular Devices). Fluorescence was excited at 480 nm and emitted light was collected above 520 nm. Fluorescence images were acquired at 1 Hz using 10% LED power and an exposure time of 400 ms. Drugs were locally applied through a gravity-fed perfusion system controlled by solenoid valves and equipped with a perfusion pencil multi-barrel manifold tip. Cells were continuously perfused with artificial cerebrospinal fluid (aCSF; 310 mOsm, pH 7.4) containing (in mM): 140 NaCl, 2 CaCl₂, 3 KCl, 10 HEPES and 10 D-glucose. All drugs were diluted in aCSF immediately before application. ATP (10 µM; A26209, Sigma-Aldrich) was first locally applied for 15 s, followed by IVM (3 µM; 18898, Sigma-Aldrich) perfusion for 3 min. Cells were subsequently exposed to a combined application of ATP and IVM for an additional 15 s. At the end of each recording, ionomycin (10 µM; HB1001, HelloBio) was applied in aCSF as a positive control to confirm GCaMP6s functionality.

Fluorescence changes were quantified from time-lapse recordings using ImageJ. Responses were expressed as the area under the curve (AUC) calculated over a 2-min period following each stimulation. Data were exported to Microsoft Excel for processing, and statistical analyses and graphical representations were performed using GraphPad Prism 8.

### Plasmid construction

Two constructs were generated using classical restriction enzyme-based cloning. The GFP coding sequence was excised from a previously available plasmid using PmeI and XbaI, and ligated into the pcDNA3.1 vector digested with the same enzymes. For the second construct, the insert containing GFP fused to the P2X4 3′UTR was released from an existing plasmid using XbaI and NotI, and subsequently cloned into the same vector backbone digested with the corresponding enzymes. In both cases, ligation products were transformed into Escherichia coli DH5α competent cells, and colonies were selected on LB agar plates containing 100 µg/mL ampicillin. Plasmid DNA was extracted using the QIAprep Spin Miniprep Kit, Qiagen, and correct insertion and orientation were confirmed by restriction enzyme digestion. ZFP36 knock-down was performed with lentiviral shRNA. pLKO encoding shRNA were obtained from Sigma (TRCN0000238325, sequence ACCACCTCCTCTCGATACAAG, and TRCN0000238326, sequence GGTCAGACTCACCTGTCTTTG). Lentiviral production was previously described [14]. Plasmids pTRIPZ Myc2-mouse TTP WT and pTRIPZ Myc2-mouse TTP aa S52 S178 MUT were gifts from Julian Downward (Addgene plasmid # 107000 and [Addgene plasmid #107001](https://www.addgene.org/107001/), respectively [15].

### Pharmacological treatment

CCL21 (40 ng/mL, 250-13, Peprotech), Actinomycin D (5 µM, HB1477, HelloBio), R18 (5 µM, BML-P214-0001, Enzo life science), SB203580 (10 µM, 152121-47-6, Sigma) were dissolved in H20, okadaic acid (1 mM in DMSO, ALX-350-003-C025, Enzo life science), were added to the culture medium for 6h.

### RNAscope in *situ* hybridization

Sections mounted on Superfrost Plus slides (Thermo Fisher Scientific) were post-fixed in 4 % PFA (1 h), dehydrated in graded ethanol (50, 70, 100 %) and baked for 1 h at 60 °C in a HybEZ™ II oven (ACD, Bio-Techne). Endogenous peroxidases were quenched in hydrogen peroxide (10 min), followed by target retrieval in boiling RNAscope® Target Retrieval solution (15 min) and RNAscope® Protease III treatment (40 °C, 30 min). Probes (*mP2rx4*, 473691; *mZfp36*, 888831; *hZFP36*, 427351; negative control, 321831; all from ACD, Bio-Techne) were pre-warmed to 40 °C and hybridized for 2 h at 40 °C. The signal was amplified with RNAscope® Multiplex FL v2 Amp1-3 (30, 30 and 15 min at 40 °C respectively), developed with RNAscope® Multiplex FL v2 HRP-C1 (15 min) and TSA® Fluorescein (30 min), and terminated with RNAscope® HRP blocker (15 min). Sections were washed in RNAscope® wash buffer between all steps and then processed for Iba1 immunofluorescence as described above.

### Image acquisition and analysis

Images were acquired with a 20× objective on an Axio Imager 2 microscope equipped with an Apotome 3 structured-illumination module (Zeiss) and a CoolSNAP HQ camera (Photometrics). Zeiss .czi files were converted to .ims with Imaris File Converter 9.9.1 (Oxford Instruments) and analyzed in Imaris 7.4.2 (Bitplane, Oxford Instruments) using the Surpass > Cells module in “Cell Body” mode. Microglia were reconstructed in 3D with smoothing and background subtraction, with a minimum object size of 330 voxels; gaps were not filled, to avoid generating aberrant processes. Reconstruction thresholds were defined on negative-probe sections and adjusted manually per image. RNAscope signal was quantified with the vesicle-detection module (estimated diameter 1 µm, background subtraction), the threshold being set so that no spot was detected on negative-probe images. Microglia were counted manually, and data are expressed as the mean number of spots per microglial cell, averaged per image and per animal.

### Statistical analysis

Data are presented as mean ± SEM and were analyzed with GraphPad Prism 8. Groups were compared using Prism software.

## Results

It is well established that P2X4 is *de novo* expressed in spinal cord microglia following spared nerve injury (SNI) [4]. To determine whether this upregulation occurs at the transcriptional level, we first examined P2X4 mRNA expression under similar conditions using fluorescent *in situ* hybridization. At 15 days post-SNI, we detected the presence of P2X4 mRNA in the dorsal horn of the spinal cord in both control and neuropathic male and female mice (Fig. 1A). Interestingly, quantification of P2X4 mRNA puncta within microglial cells showed comparable levels between control and SNI groups (Fig. 1B), suggesting that the increased P2X4 protein expression is not driven by increased transcription. To further investigate this hypothesis, we performed ddPCR on dorsal horn of the spinal cord samples from *Cx3Cr1*^CreERT2/+^;*Rpl22*^HA(RiboTag)/+^ mice (Fig. 1C). In contrast to the RNAscope data, ddPCR analysis revealed a significant increase of ribosome associated P2X4 mRNA in the ipsilateral side of the lesion following SNI in both sexes compared to the contralateral dorsal horn (male: sham ipsi: 0.092 ± 0.058; sham contra: 0.105 ± 0.03; SNI ipsi: 0.405 ±0.04; SNI contra: 0.1 ± 0.03. Female: sham ipsi: 0.18 ± 0.03; sham contra: 0.183 ± 0.05; SNI ipsi: 0.303 ± 0.005; SNI contra: 0.183 ± 0.02) indicating an increase of mRNA translation. To confirm this translational effect, we treated BMDM cultures with CCL21, a chemokine known to induce P2X4 upregulation [16], for 6 hours and assessed P2X4 protein levels by western blot. CCL21 treatment led to a significant increase of P2X4 protein (204.2 ± 32.39 %) relative to normalized basal conditions (Fig. 1D). This upregulation was not prevented by the co-application of Actinomycin D (283.9 ± 32.82 %), a transcription inhibitor. Together, these results support a post-transcriptional mechanism of P2X4 regulation.

**Figure 1:**
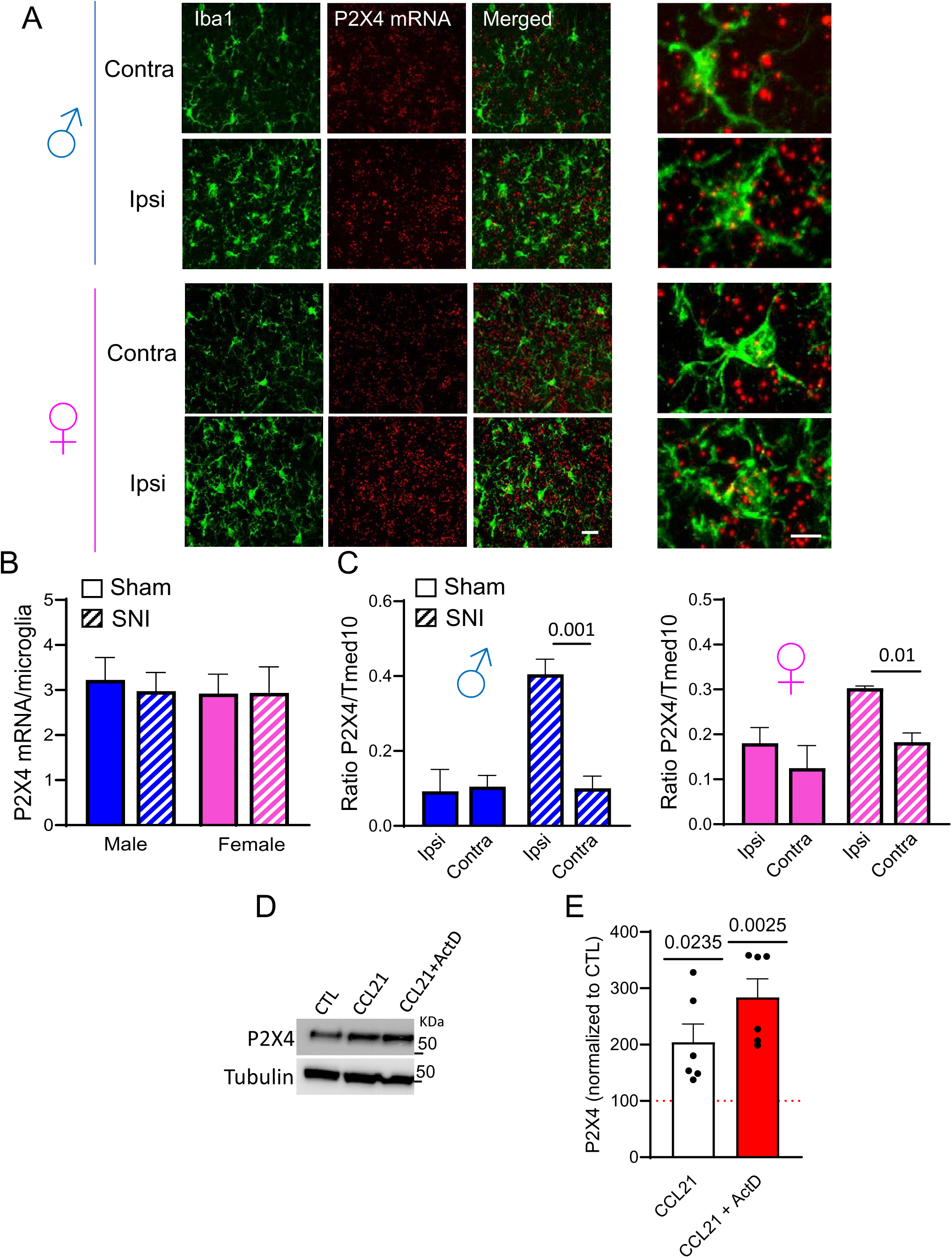
Microglial P2X4 mRNA is translated during neuropathy in the dorsal horn of the spinal cord. **(A)** Representative images of the dorsal horn of the spinal cord in sham and neuropathic mice, on the ipsilateral and contralateral sides in male and female (15 days post neuropathy). Immunohistochemical staining of microglia (Iba1, green) and RNAscope staining of P2X4 mRNA (red). Scale bar = 20 μm. **(B)** Quantification of the mean number of P2X4 mRNA molecules per microglia in the ipsilateral dorsal horn of male and female sham and SNI mice. Microglia express equivalent levels of P2X4 mRNA under sham and SNI conditions in both sexes. Mean ± SEM; n = 11-13 sections from 3 different animals; two-way ANOVA. **(C)** Quantification by ddPCR of P2X4 mRNA copy numbers in spinal microglia in sham and SNI animals after 15 days, normalised to the house gene Tmed10. The results show an increase in translating mRNA in ipsilateral microglia of the dorsal horn in neuropathic conditions compared with the contralateral side and sham animals in male and female mice. Mean ± SEM, n = 3-4, each data point represents 4-6 animals. two-way ANOVA. **(D)** Representative western blot of P2X4 in BMDM cells treated with CCL21 (40 ng/mL) ± Actinomycin D (ActD, 5 µM). **(E)** Quantification of P2X4 expression detected in (D), normalized to basal condition. CCL21 induces P2X4 increase that is not prevented by ActD. N=6 independent experiments, mean ± SEM, one sample t-test.

To gain insight into the post-transcriptional mechanisms modulating P2X4 translation, we focused on the 3′UTR sequence that often carries miRNA and RNA-binding protein (RBP) regulatory sequences. To this end, we generated a luciferase reporter construct containing or not the 1424 bases full-length P2X4 3′UTR (Fig. 2A). These constructs were transfected into HEK293 cells and reporter activity was measured. As shown in Fig. 2B, C, transfection of the P2X4 3′UTR-containing construct led to a significant decrease in reporter activity compared to cells transfected without P2X4 3′UTR (170.9 ± 27.65 vs 488.6 ± 39.87). To further validate these findings at the protein level, we performed western blot analysis on HEK293 cells transfected with P2X4 with or without the 3’UTR sequence constructs. As presented in Fig. 2D, E, protein expression from the P2X4 3′UTR-containing transfected construct was markedly reduced (71.58 ± 6.5 %) compared to the normalized control, confirming that the 3′UTR negatively regulates P2X4 protein levels. These results suggest that 3′UTR-mediated regulation fine-tunes P2X4 protein levels independently of transcriptional control.

**Figure 2:**
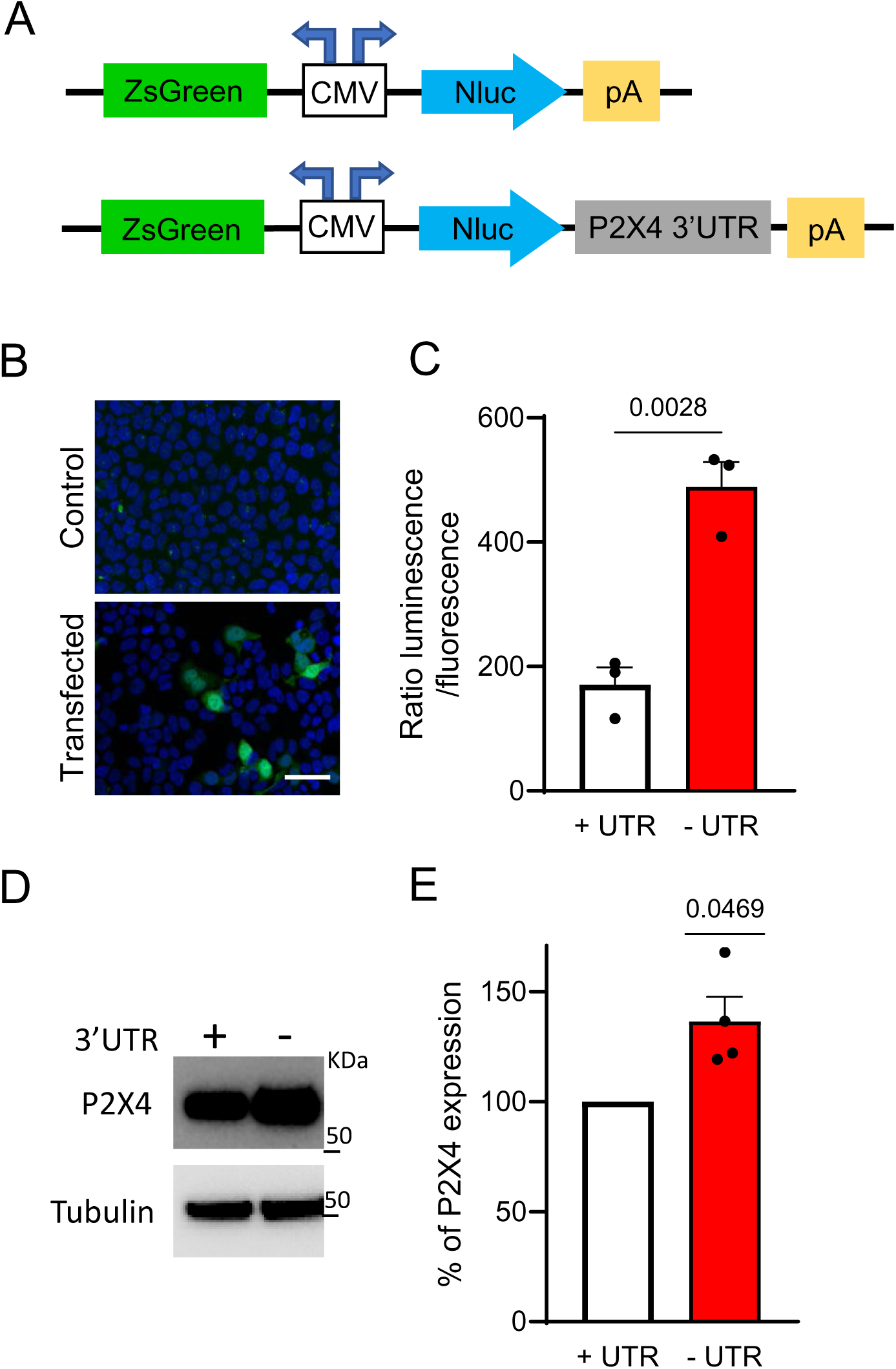
P2X4 3′UTR mRNA sequence regulates P2X4 expression in HEK293 cells. **(A)** Schematic representation of mP2X4-GFP ± 3′UTR plasmid constructs. **(B)** Representative image of HEK cells transfected or not with mP2X4-GFP plasmid. Green: GFP; Blue: DAPI. Scale bars: 30 µm. **(C)** Quantification of the luminescence normalized to the GFP fluorescence detected in (B). The absence of 3′UTR of P2X4mRNA induces a significant increase of the ratio. N= 3 independent experiments, mean ± SEM, Student t test. **(D)** HEK cells were transfected with plasmids encoding P2X4 either with (+3′UTR) or without (−3′UTR) its mRNA 3′ untranslated region. P2X4 protein expression was assessed by Western blot. **(E)** Densitometric quantification of P2X4 expression obtained in (D). N= 3 independent experiments, mean ± SEM, one sample t-test.

Having established that the 3′UTR of P2X4 represses protein expression, we next asked whether this effect could be mediated by RNA-binding proteins (RBPs), which are known to modulate mRNA stability and translation through interaction with 3′UTR regulatory elements. Using bioinformatic prediction tools, we identified ZFP36 as a potential candidate RBP that could interacts with the P2X4 3′UTR through ARE binding motifs (Fig. 3A). To determine the involvement of ZFP36 in the control of P2X4 expression, we transfected HEK293 cells with constructs encoding either wild-type (WT) ZFP36 or a phosphorylation-deficient mutant of ZFP36 in which key phosphorylation sites were mutated [17]. As shown in Fig. 3B, C, expression of WT ZFP36 significantly reduced P2X4 protein levels. In contrast, the phosphorylation-deficient mutant induced a significant increase of P2X4 expression, indicating that phosphorylation of ZFP36 is required for its repressive function on P2X4 mRNA. To further dissect the signaling pathways that modulate ZFP36-dependent activity, we transfected HEK293 cells with the P2X4 3′UTR luciferase reporter construct and treated them with pharmacological modulators targeting pathways known to regulate ZFP36 phosphorylation and function. As shown in Fig. 3D, treatment with SB203580 (+SB), an inhibitor of p38 MAPK, significantly decreased reporter luminescence (76.72 ± 7.51 %) compared to control conditions. Similarly, inhibition of the ZFP36 chaperon 14-3-3 with R18 (+R18, a competitive inhibitor of 14-3-3) also led to a significant reduction in reporter activity (81.49 ± 5.34 %). These results suggest that both p38 MAPK and 14-3-3 signaling pathways modulate P2X4 expression through its 3′UTR, potentially by regulating ZFP36 phosphorylation and activity. Treatment with okadaic acid (+OA), a phosphatase inhibitor, significantly increased reporter activity (150.9 ± 14.39 %), further supporting the involvement of phosphorylation-dependent signaling cascades in the regulation of P2X4 by endogenous ZFP36.

**Figure 3:**
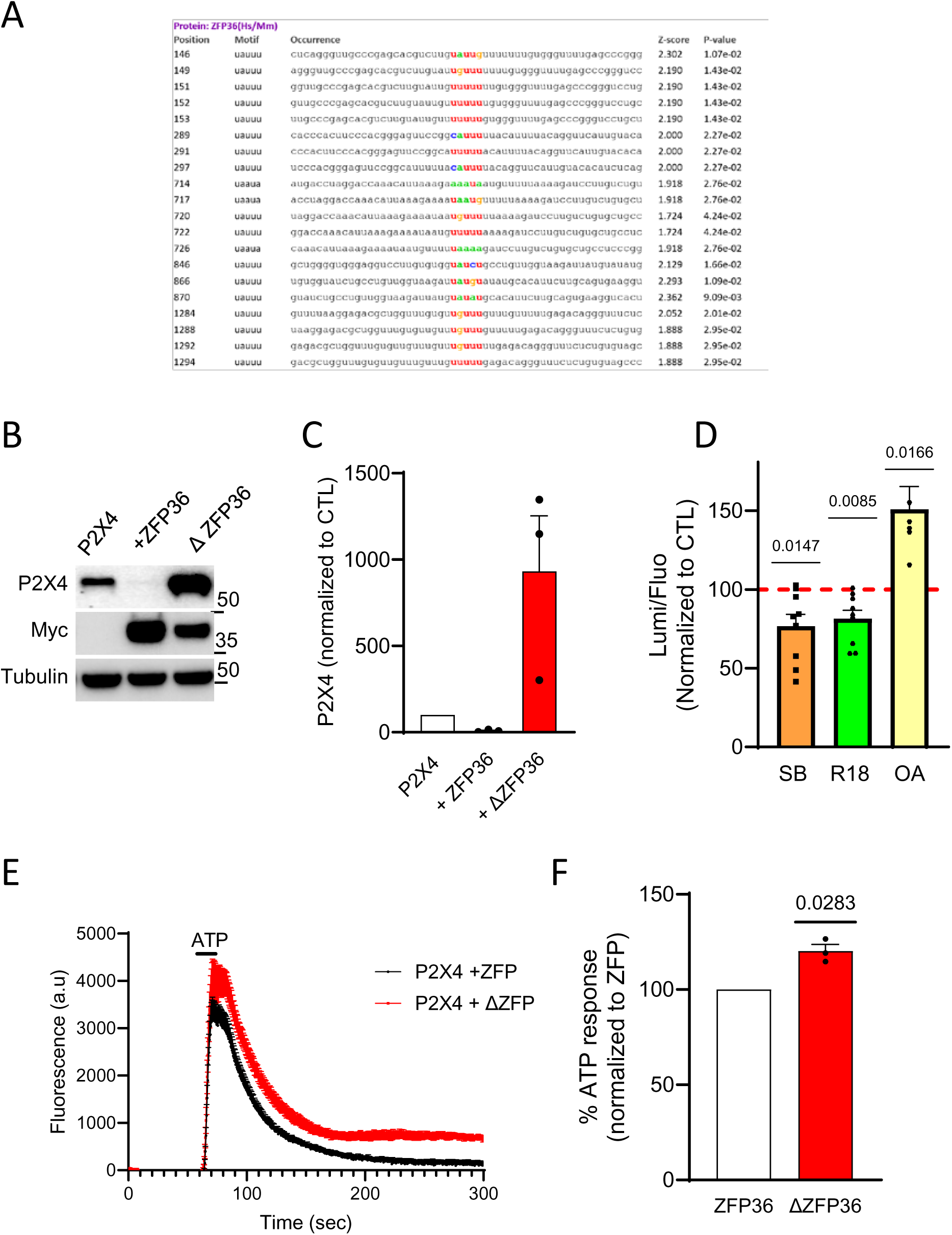
ZFP36 controls P2X4 mRNA translation. **(A) Potential ZFP36-binding sites within the mRNA of P2X4.** Prediction of potential ZFP36-binding sites along the P2X4 mRNA sequence. For each predicted site, the position, recognized motif, nucleotide sequence context Z-score and associated *p*-value are shown. Colored nucleotides highlight the potential ZFP36-binding motifs within the RNA sequence. **(B)** Representative western blot of HEK293 cells transfected with P2X4 and ZFP36 or mutated ZFP36 (ΔZFP36). **(C)** Quantification of P2X4 expression obtained in (B), normalized to control condition. The presence of mutated ZFP36 induces a significant increase of P2X4 expression. N= 3 independent experiments, mean ± SEM, Student t test **(D)** Quantification of the luminescence normalized to the GFP fluorescence detected in HEK293 cells transfected with P2X4 cDNA, normalized to control conditions. The inhibition of p38 MAPK (SB) or 14-3-3 (R18) induces a significant decrease of P2X4 expression. The inhibition of phosphatase 1 inhibitor (okadaic acid, OA) increases P2X4 expression. N= 8 independent experiments, one sample t-test. **(E)** HEK293 cells were co-transfected with P2X4 together with either wild-type (WT) ZFP36 or a phosphorylation-deficient mutant of ZFP36 and stimulated with ATP (20 µM). Representative calcium traces showing the kinetic response to ATP in cells expressing P2X4 with WT ZFP36 (black) or with the ZFP36 mutant (red). **(F)** Quantification of the peak amplitude of the calcium response. n=3 independent experiments mean ± SEM, unpaired t-test. Cells expressing the phosphorylation-deficient ZFP36 mutant displayed a significantly increased calcium response compared to WT ZFP36-expressing cells.

Finally, to confirm that ZFP36-mediated regulation of P2X4 expression has functional consequences, we transfected HEK293 cells with P2X4 cDNA together with either WT ZFP36 or the phosphorylation-deficient mutant of ZFP36. Intracellular calcium measurements were performed using the FDSS plate reader following ATP stimulation (20 µM). As shown in Fig. 3E, cells co-expressing P2X4 and WT ZFP36 displayed a smaller calcium response upon ATP application compared to cells co-expressing P2X4 and phosphorylation-deficient ZFP36 (120.2 ± 3.46 %, Fig. 3F). Of note, the residual ATP-induced calcium response observed in the presence of WT ZFP36 is certainly attributable to endogenous P2Y receptor activation. Altogether, these results provide functional evidence that ZFP36 modulates P2X4 expression, and consequently P2X4 channel activity, through mRNA/3′UTR interaction.

To determine whether ZFP36 regulates endogenous P2X4 expression in a non-recombinant cellular context, we knocked down ZFP36 in primary macrophages (BMDMs) and assessed P2X4 expression. BMDMs were transduced either with shRNA expressing lentivirus targeting ZFP36 or with a GFP-expressing virus. As shown in Fig. 4A, B, knockdown of ZFP36 induced a significant increase of P2X4 protein levels (846.4 ± 196.5 %) compared to normalized control cells, suggesting that ZFP36 represses P2X4 expression under basal conditions.

**Figure 4:**
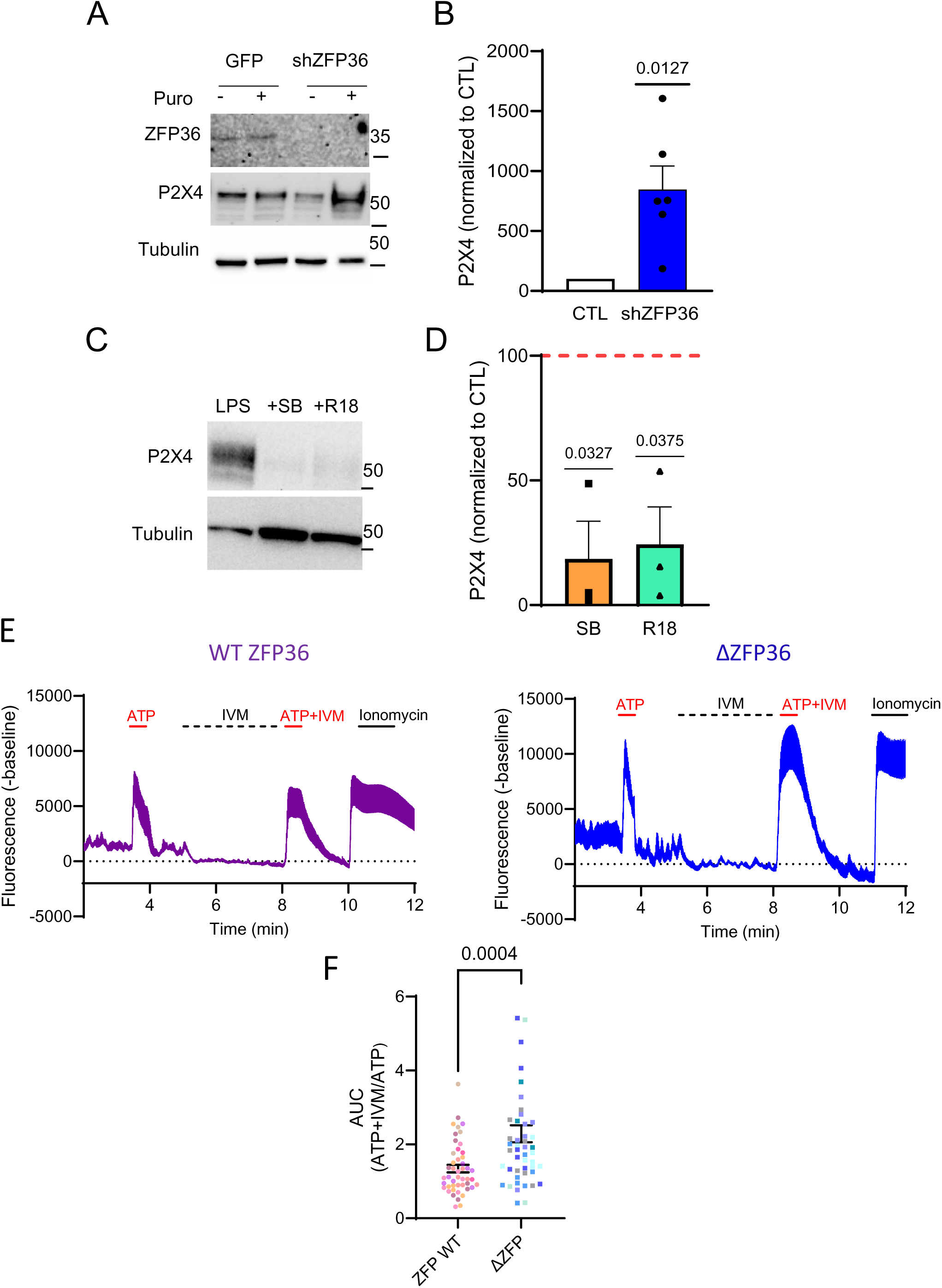
ZFP36 controls P2X4 expression in BMDM cells. **(A)** Representative western blot of BMDM cells transduced with control (GFP) or ZFP36 shRNA lentivirus. **(B)** Quantification of P2X4 expression obtained in (A), normalized to control condition. The inhibition of ZFP36 induces a significant increase of P2X4 expression. N= 6 independent experiments, one sample t-test. **(C)** Representative western blot of BMDM cells treated with p38 MAPK (SB) or 14-3-3 (R18) inhibitor. **(D)** Quantification of P2X4 expression observed in (C). Both inhibitors decrease P2X4 expression. N= 3 independent experiments, one sample t-test. **(E)** Representative recordings of ATP-induced calcium signals in individual macrophages expressing ZFP36 (left) or ZFP36-mut (right), together with cytosolic GCaMP6s. ATP (10 µM) was locally applied for 15 s, followed by ivermectin (IVM, 3 µM) for 3 min. Cells were subsequently stimulated with ATP (10 µM) in the continued presence of IVM for an additional 15 s. **(F)** Population analysis of ATP-induced calcium responses. The area under the curve (AUC) was calculated over a 2-min period following each stimulation. The AUC of the ATP + IVM response was normalized to that of the initial ATP application in the absence of IVM. Each point represents an individual cell, and cells from the same recording are shown in the same color. Data were obtained from >47 cells, unpaired t-test.

To further confirm the role of ZFP36, we treated BMDMs with SB203580 or R18 and assessed P2X4 expression by western blot. As shown in Fig. 4C, D, treatment with either inhibitor led to a decrease of P2X4 protein levels compared to untreated cells (SB: 18.53 ±15.11 %, R18: 24.34 ± 15.07 %) consistent with the reporter assay results obtained in HEK293 cells.

We next investigated whether the functional activity of endogenous P2X4 could be modulated by ZFP36. Cells were transduced using lentivirus expressing GCamp6 and lentivirus expressing either WT or mutated ZFP36. Because ATP activates both P2X and P2Y receptors, P2X4 contribution to global calcium signal was revealed through the use of ivermectin, a positive allosteric modulator of P2X4. ATP stimulation of WT ZFP and phosphorylation-deficient mutated ZFP transduced BMDM induced a transient increase in fluorescence in both conditions (Fig. 4E, F). However, following ivermectin (IVM) treatment, a positive allosteric modulator of P2X4, subsequent ATP stimulation (ATP + IVM) elicited a significant increase of fluorescence response in mutant ZFP36-expressing BMDMs compared to WT ZFP36**-** expressing cells. Consistent with these representative traces, quantitative analysis of the area under the curve (AUC) for the ATP + IVM response, normalized to the response to ATP alone, revealed a significant increase in mutant ZFP36-expressing BMDMs (2.288 ± 0.22) compared with WT ZFP36-expressing BMDMs (1.344 ± 0.1). These results indicate that loss of functional ZFP36 is associated with an enhanced activity of P2X4.

Finally, we examined whether this regulatory mechanism operates in primary microglial cells. Immunocytochemical analysis of primary microglial cultures revealed that LPS stimulation induced a marked increase in P2X4 immunoreactivity (Fig. 5A). Interestingly, this LPS**-**induced increase was inhibited upon co**-**treatment with SB203580 or R18, further supporting the involvement of p38 MAPK and 14-3-3 signaling pathways in the regulation of P2X4 expression in microglia. Collectively, these findings indicate that ZFP36 negatively regulates P2X4 expression in a phosphorylation-dependent manner and that this regulation is controlled by signaling pathways involving p38 MAPK, 14-3-3 and phosphatases that converge to modulate ZFP36 activity. To confirm that ZFP36 is expressed in spinal cord microglia, we performed RNAscope *in situ* hybridization on lumbar spinal cord sections from naive male mice. As shown in Fig. 5B, ZFP36 mRNA was detected, but not exclusively, in Iba1-positive cells, indicating that ZFP36 is expressed in spinal microglia.

**Figure 5:**
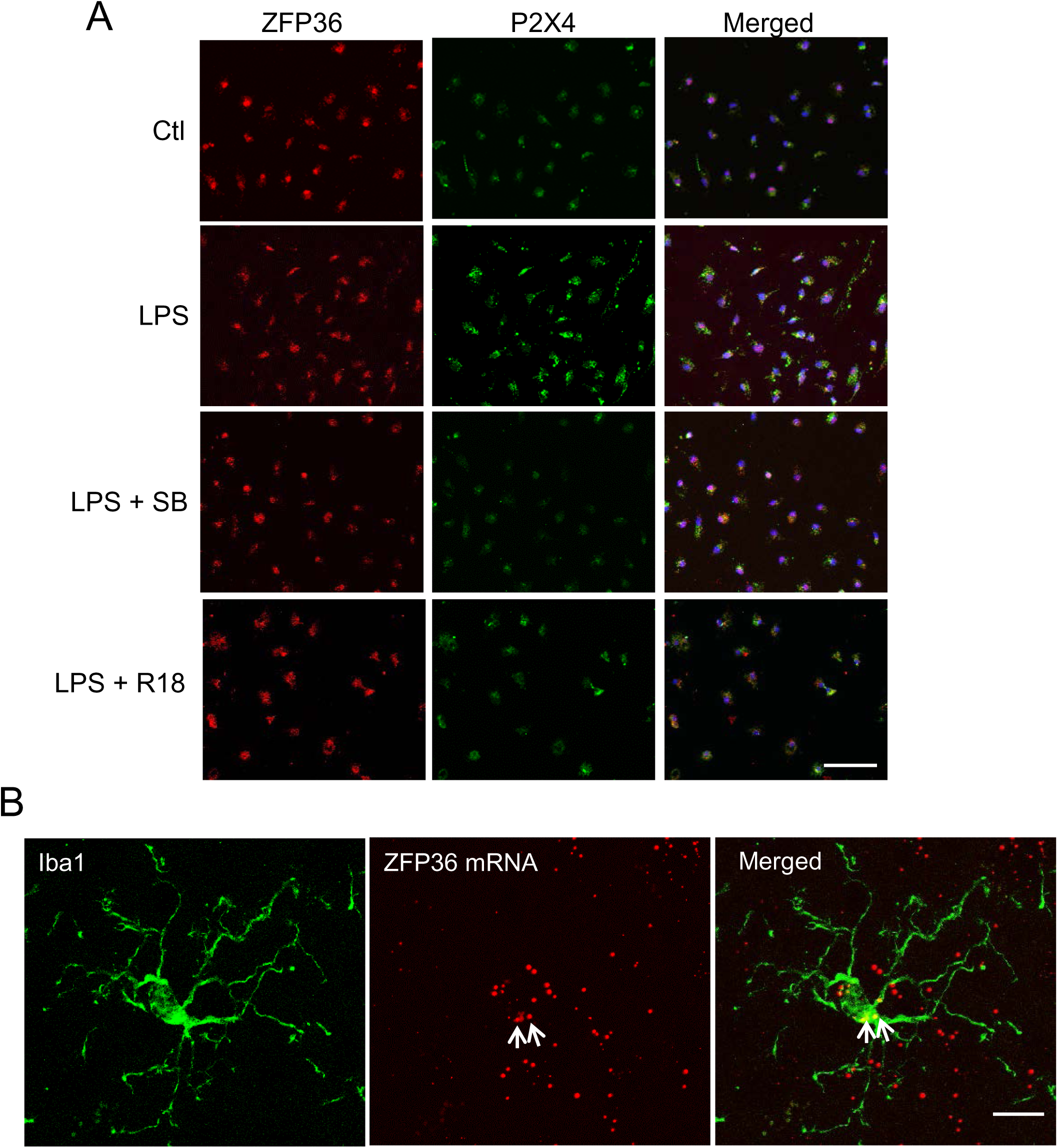
ZFP36-associated signaling regulates P2X4 expression in microglia. **(A)** Representative immunocytochemistry images of primary microglial cultures treated with LPS alone or in combination with the p38 MAPK inhibitor SB203580 (SB) or the 14-3-3 inhibitor R18. P2X4 immunoreactivity (green) was increased by LPS and reduced by co-treatment with SB or R18. Scale bar: 50 µm. **(B)** Representative RNAscope images showing ZFP36 mRNA (red) in Iba1-positive microglia (green) in the dorsal horn of the mouse spinal cord. Scale bar: 10 µm.

Altogether, these results provide functional evidence that ZFP36 inhibits P2X4 translation, and consequently P2X4 channel activity, through mRNA/3′UTR regulation in macrophages and microglia.

## Discussion

In the present study, we identify the RNA-binding protein ZFP36 as a novel post-transcriptional regulator of P2X4 expression in microglia and macrophages. Our data show that ZFP36 represses P2X4 protein expression through the mRNA 3′UTR, which contains predicted AU-rich elements (AREs) and that this regulation is phosphorylation-dependent, involving p38 MAPK, 14-3-3, and phosphatase signaling, known to regulate the RNA-binding activity of ZFP36 [17]. Given the established role of P2X4 in both inflammatory and neuropathic pain [3] [4] [18], this mechanism may shape microglial purinergic responsiveness after nerve injury and represents a potential therapeutic target.

ZFP36 is known for destabilizing cytokine mRNAs such as TNF-α and IL-6 [19] [7] [20] [21], and its genetic deletion in mice causes a severe systemic inflammatory syndrome [8]. Interestingly, our work extends its repertoire to a purinergic receptor. While P2X4 protein is strongly induced in microglia following SNI [4], our RNAscope analysis revealed that microglial P2X4 mRNA levels remain unchanged. Yet, RiboTag-based ddPCR showed increased ribosome-associated P2X4 mRNA on the ipsilateral side of nerve lesion. This dissociation between total and translating mRNA supports a post-transcriptional mechanism contributing to P2X4 upregulation, complementing the transcriptional IRF8–IRF5 axis previously described [22]. Together, these observations suggest that transcriptional and post-transcriptional mechanisms cooperate to achieve the full upregulation of P2X4 protein in neuropathic conditions.

Mechanistically, mutation of key phosphorylation sites abolished ZFP36-mediated repression and pharmacological modulation of p38 MAPK, 14-3-3 and phosphatases consistently altered P2X4 expression through its 3′UTR in macrophages and microglia. These results are consistent with the model in which ZFP36 phosphorylation controls its RNA-binding activity and subcellular localization [17][23] [24] [25], thereby tuning P2X4 mRNA translation efficiency. They also align with the known regulation of TTP-family proteins by p38/MK2 and 14-3-3 and with the involvement of p38 in spinal microglia after nerve injury [24] [25]. Notably, p38 inhibition or 14-3-3 disruption decreased P2X4 expression, whereas phosphatase inhibition increased it, consistent with a phosphorylation-dependent switch controlling ZFP36 activity.

Our findings intersect with several independent lines of evidence. First, the post-transcriptional control of ion channels and receptors by ARE-binding proteins is increasingly recognized as a mechanism shaping neuronal and glial excitability. For instance, ZFP36 has been shown to regulate the stability of mRNAs encoding inflammatory mediators in microglia [26] and other RBPs such as HuR and TIA-1 modulate the translation of pain-related transcripts [27] [28]. Our study extends this concept to the ligand-gated ion channel P2X4, establishing a direct link between RNA metabolism and purinergic signaling in the spinal cord.

The interplay between transcriptional and post-transcriptional control observed here extends previous work on P2X4 regulation, which had primarily focused on transcriptional mechanisms. The IRF8–IRF5 axis drives P2X4 transcription after nerve injury [22] but the absence of a corresponding increase in total P2X4 mRNA despite robust protein induction implies that additional layers of control exist [4]. Our RiboTag data provide direct evidence for such a layer, showing that translating P2X4 mRNA increases without changes in total mRNA. This is consistent with observations in other systems where translational efficiency, rather than mRNA abundance, is a major determinant of protein output [29].

The involvement of p38 MAPK in our mechanism is particularly relevant to neuropathic pain. p38 is activated in spinal microglia after nerve injury and is a well-established contributor to microglial-mediated pain hypersensitivity [4] [30] [31]. Our data suggest that p38 may exert part of its pro-nociceptive effects through ZFP36-dependent stabilization of P2X4 translation. This suggests that ZFP36-P2X4 axis downstream of a signaling pathway already validated as a therapeutic target in pain.

Finally, our results align with the broader concept of “post-transcriptional operons” in inflammation, where coordinated regulation of mRNA stability and translation allows rapid and reversible control of gene expression [32]. ZFP36, as a key player, may thus coordinate the expression of both cytokines and purinergic receptors, ensuring that microglia integrate inflammatory and purinergic cues during neuropathic pain.

The ZFP36-P2X4 axis may have implications beyond P2X4 itself. Microglia express multiple purinergic receptors, including P2X7, P2Y6 and P2Y12, several of which are post-transcriptionally regulated through cis-acting elements in their 3′UTRs [33] [34] [35]. It is tempting to speculate that ZFP36 may coordinate the expression of a broader purinergic module, thereby tuning microglial sensitivity to extracellular ATP. In support of this idea, ZFP36 deficiency in mice leads to exacerbated inflammatory responses [8] associated with the role of ATP as a key danger signal in inflamed and injured tissues [36]. Thus, ZFP36 may serve as a checkpoint that restrains purinergic signaling under basal conditions while permitting injury-induced P2X4 upregulation. Moreover, the regulation of P2X4 by ZFP36 in both macrophages and microglia suggests that this mechanism may operate in peripheral immune cells as well. P2X4 is expressed in macrophages [37] and contributes to inflammasome activation and cytokine release by potentiating P2X7-dependent signaling [37] [38]. Future studies should explore whether the ZFP36-P2X4 axis is conserved across myeloid cell types and disease contexts.

However, whether ZFP36 directly binds P2X4 mRNA remains to be established. Similarly, we cannot exclude that p38 and 14-3-3 also act through other RBPs [39] or that additional regulatory elements within the P2X4 3′UTR contribute to this regulation. Finally, although P2X4 mRNA translation increases similarly in male and female neuropathic mice, sex differences in microglial P2X4 signaling have been reported in other contexts [40] [41] [42] and whether the ZFP36-P2X4 axis is sexually dimorphic remains an open question.

Collectively, our findings uncover a phosphorylation-dependent, post-transcriptional mechanism by which ZFP36 controls P2X4 expression in microglia and macrophages, extending the functional repertoire of ZFP36 beyond cytokine regulation to a purinergic receptor. This ZFP36-P2X4 axis may act as a checkpoint controlling microglial responsiveness to extracellular ATP, thereby shaping the sensitivity of myeloid cells to purinergic signals under both physiological and pathological conditions. Given the established role of P2X4 in neuropathic pain, targeting mechanisms involved in the regulation of P2X4 expression may represent a potential pathway for future therapeutic intervention in chronic pain.

## Acknowledgements

We thank the following Montpellier Biocampus core facilities: Réseau des Animaleries de Montpellier (RAM) (Iexplore facility), the imaging facility MRI, *Réseau d’Histologie Expérimentale de Montpellier (RHEM) and ARPEGE facility*.

## Author contributions

D.G, N.L, N.A, L.G, V.C L.U, F.R performed the experiments and analyzed the data. F.R and L.U conceived the study and designed the experiments. L.U wrote the manuscript. All authors commented on the manuscript.

## Availability of data and materials

All data generated or analyzed during this study are included in this published article.

## Declarations

Ethics approval and consent to participate: All experiments followed European Union (Council directive 2010/63/UE) and institutional guidelines for the care and use of laboratory animals. The animal experiment protocols used in this study were approved by the Comité d’Ethique pour l’Expérimentation Animale Languedoc Roussillon (CEEA-LR; APAFiS#22741).

## Consent for publication

Not applicable.

